# Integrative multi-modal analysis reveals the contribution of noncoding RNAs to post-treatment progression of IDH-mutant astrocytomas

**DOI:** 10.1101/2025.09.30.679694

**Authors:** Anja Hartewig, Serafiina Jaatinen, Matti Annala, Joonas Haapasalo, Wei Zhang, Hannu Haapasalo, Matti Nykter, Xin Lai, Kirsi J. Rautajoki

## Abstract

IDH-mutant (IDHmut) astrocytomas typically arise as grade 2–3 tumors, but a subset of them progress to grade 4 after treatment, significantly worsening the prognosis. It is unresolved how noncoding RNAs (ncRNAs) contribute to this tumor progression. To characterize ncRNAs’ regulatory roles, we profiled and analyzed the coding and noncoding transcriptomes of matched tumor samples before and after progression to grade 4 in IDHmut astrocytoma patients. By integrating our data with public primary and matched tumor cohorts, we found that upregulated protein-coding genes in progressed tumors overlapped with those in primary grade 4 tumors and were linked to cell proliferation. In contrast, downregulated genes differed between primary and post-treatment grade 4 tumors. A large fraction of genes that were downregulated only in the post-treatment setting were associated with decreased cell differentiation. We identified 53 progression-related ncRNAs predicted to regulate 125 differentially expressed genes. Gene regulatory network analysis revealed their involvement in cell cycle control, extracellular matrix (ECM) organization, and neural differentiation. Notably, hsa-let-7b-3p, a tumor suppressor microRNA, showed recurrent hemizygous deletions and downregulation after post-therapy progression. The long noncoding RNA (lncRNA) PVT1 was recurrently gained and upregulated in grade 4 tumors. The lncRNA NEAT1, previously linked to treatment resistance, was especially upregulated post-treatment and had the highest number of predicted targets (n=38), many related to ECM organization. Overall, our findings highlight the role of ncRNAs in the post-treatment progression of IDHmut astrocytomas, offering new insights into mechanisms of their malignancy.

## Introduction

Despite accounting for <2% of newly diagnosed cancer cases annually, diffuse astrocytomas are associated with substantial mortality and morbidity [1]. The median age at diagnosis is 68–70 years for IDH wild-type (IDHwt) glioblastoma (GB) (the most aggressive subtype of diffuse astrocytoma) [2]. By contrast, IDH-mutant (IDHmut) astrocytomas are typically diagnosed in younger adults, with a median age of 36 years at diagnosis of grade 2–3 tumors [3]. The median overall survival (OS) for grade 2–3 IDHmut astrocytomas is 7–10 years [4]. Many tumors relapse and ultimately progress to grade 4, significantly worsening the prognosis.

The initial treatment for grade 2–3 gliomas involves maximal safe resection. Depending on the extent of the residual tumor after surgery and other risk factors, adjuvant treatments, such as radiation and chemotherapy, can be administered [5]. Although adjuvant treatment prolongs survival, it may also contribute to tumor evolution. For example, radiation increases the risk of homozygous *CDKN2A* deletion [6,7], an alteration frequently observed in recurrent grade 4 IDHmut astrocytomas [7,8]. The DNA alkylating agent temozolomide (TMZ) can induce a hypermutated phenotype [9]. The combination of young age and treatment side effects, such as cognitive impairment [10,11], and the potential influence of treatment on tumor evolution [12,13], highlights the need for better treatment options and predictive biomarkers.

Noncoding RNAs (ncRNAs) are functional RNA molecules. These include microRNAs (miRNAs), which are 18–25 nucleotide long, single-stranded molecules, and long noncoding RNAs (lncRNAs), which are longer than 200 nucleotides [14]. miRNAs post-transcriptionally regulate the expression of protein-coding genes [15]. LncRNAs regulate transcription and are involved in chromatin remodeling and protein regulation [16]. The competing endogenous RNA (ceRNA) hypothesis proposes that lncRNAs and mRNAs compete for miRNA binding because they contain similar miRNA response elements, thereby forming complex regulatory networks [17]. Both miRNAs and lncRNAs have been implicated in GB initiation [18,19], cell proliferation [20,21], and treatment resistance [22,23]. Gene regulatory networks (GRNs) have emerged as powerful models for unraveling the combinatorial functions and roles of ncRNAs under different pathological conditions [24,25]. Furthermore, ncRNAs serve as promising diagnostic and prognostic biomarkers and predictors of treatment responses. For example, miR-210 expression in gliomas is induced by hypoxia [26,27], can be measured in serum, and serves as a prognostic factor for glioma patient survival [28]. In addition, serum lncRNA MALAT1 expression is a predictive marker of TMZ treatment response in GB [29].

Despite recent advances, the role of ncRNAs in IDHmut astrocytoma progression, particularly in the context of adjuvant treatment, remains to be elucidated. Furthermore, the rarity of these entities and their potentially long time-to-progression pose challenges in obtaining matched samples for detailed analysis. In this single-center study, we investigated the role of ncRNAs (miRNAs and lncRNAs) in the post-treatment progression of diffuse IDHmut astrocytomas from grades 2 and 3 to grade 4. We investigated the changes in gene expression, characterized dysregulated ncRNAs with respect to their targets and DNA copy number alterations, and constructed a GRN. Findings related to protein-coding genes were validated using both the GLASS cohort, which contains matched cases with treatment information, and TCGA primary tumor cohort. TCGA miRNA and lncRNA expression data were used, revealing both genes associated with a higher grade as well as distinct, progression-related genes.

## Materials and methods

### Study cohorts

Matched brain tumor samples obtained before and after progressing to grade 4 were collected for this study from six patients. An experienced neuropathologist evaluated FFPE tumor samples and determined the histopathological type and grade according to the criteria presented by the WHO [30]. Frozen tumor sections were stained with hematoxylin and eosin to estimate the tumor content in the samples before DNA and RNA isolation. When frozen tumor samples were sectioned for isolation, every third section was used for RNA isolation, and the others were used for DNA isolation.

The data of the TG discovery cohort was complemented with data from TCGA and GLASS consortia. TCGA data includes mRNA, lncRNA, and miRNA measurements from up to 226 primary cases (relapse samples have been excluded), as well as CNA data from 712 primary glioma cases. The GLASS consortium provides mRNA data from eight primary grade 2–3 tumors with matched data after progression to grade 4, including treatment information and CNA data. Additional matched relapse tumors were also included in the analysis of CNAs in the GLASS cohort from the same case when available.

### Analysis of TG data

#### RNA sequencing

RNA was isolated with the mirVana™ miRNA Isolation Kit (Life Technologies) so that small (<200 nucleotides) and large RNA (>200 nucleotides) molecules were divided into separate fractions. The library construction and sequencing of the RNA samples were performed at BGI using Illumina HiSeq 2000 technology for sequencing. Transcriptome sequencing was prepared for the large RNAs with a strand-specific 140-160 bp short insert library protocol, and at least 8 Gb of clean paired-end (PE90) data was obtained per library. Small RNAs were prepared for small RNA sequencing with TruSeq Small RNA Library Prep Kit (Illumina), and on average 20 Mb of clean single-end (SE50) data was obtained per library.

#### Identification of differentially expressed genes and CNAs

Differentially expressed genes were calculated as described previously [7]. Briefly, RNA sequencing reads were aligned with STAR version 2.7.0 against the human genome assembly GRCh38 using GENCODE version 30 annotations. Gene expression was counted using the featureCounts algorithm via the Rsubread package version 1.34.6. The quality of the data was controlled with a correlation plot (Fig. S1b). Read count normalization and paired differential expression between the genes before and after progression (Grade 2 and 3 vs. 4) were calculated using DESeq2. Genes with an adjusted p-value below 0.05 (BH method) and |log2foldchange| > 1 were considered significant. Copy number analysis results from cases TG01 - TG05 were obtained as described previously [7]. Briefly, the number of aligned sequencing reads was counted for each 1000 bp genome window tile. Coverage log ratios were calculated relative to a normal kidney tissue sample WGS data, median- decimated 200-fold and exported as Integrative Genomics Viewer (IGV) tracks for visual interpretation. For TG06, only targeted DNA-seq was available and it was not used for the analysis of ncRNA CNAs.

#### Identification of differentially expressed lncRNAs

Annotations for long noncoding RNA genes (version GRCh38.p12) were downloaded from Gencodegenes (https://www.gencodegenes.org/human/release_30.html). Within the obtained DE results, lncRNAs with an adjusted p-value below 0.05 (BH method) and |log2foldchange| > 1 were considered differentially expressed.

#### LncRNA target identification

Targets were downloaded from NcPath [31] and LncTarD 2.0 [32] for each DE lncRNA. Similarly as for miRNA, both Spearman’s and Pearson’s correlation coefficients were used for filtering to account for the small cohort size and high inter-patient heterogeneity. The targets were filtered to contain only correlating targets (|Spearman’s correlation coefficient| > 0.5, with a p-value < 0.05 and |Pearson’s correlation coefficient| > 0.5) and the correlating targets were filtered for significance (adj. p-value (BH method) < 0.05).

#### Identification of differentially expressed miRNAs

Mature miRNA sequences were acquired from miRBase version 22 [33], then five isomiRs were generated for each miRNA by either adding a single nucleotide (+A, +C, +G, or +T) or by removing the terminal nucleotide (-N). Sequencing reads were extracted from FASTQ files and 3’ adapters were trimmed. Adapter-trimmed reads were aligned against the catalog of miRNA and isomiR sequences (only perfect matches were accepted), producing a miRNA read count matrix. IsomiR read counts were pooled for each miRBase v.22 annotated miRNA. The quality of the data was controlled with a correlation plot (Fig. S1a). Using DESeq2, read counts were normalized using the median of ratios method [34] and differential expressions for matched tumor samples (Grade 2 and 3 vs. 4) were calculated. miRNAs with an adjusted p-value (BH method) below 0.05, an |log2foldchange| > 1 and a raw median read count of at least 10 in grade 2-3 as well as grade 4 samples were considered differentially expressed.

#### Prediction of miRNA targets and enrichment analysis of targets

For the differentially expressed miRNAs, we acquired target predictions from miRWalk [35] database, requiring that the target was predicted by at least two of the following algorithms: miRWalk 3.0, TargetScan 8.0 and miRDB 6.0, or that the target was validated experimentally in mirTarBase 7.0 database. The predictions were further filtered by using the expression correlation between miRNA and their target genes in our cohort. To account for the small cohort size and high inter-patient heterogeneity, both Spearman’s and Pearson’s correlation coefficients were used for filtered. Targets were considered if the Spearman’s correlation coefficient was below -0.5, with a p-value < 0.05 and the Pearson’s correlation coefficient was below -0.5. Additionally, only differentially expressed targets (adj. p-value (Benjamini-Hochberg (BH) method < 0.05) were considered significant for further analysis.

#### Identification of enriched Reactome pathways

The enrichR package version 3.2 was utilized to calculate significant pathways, selecting the “Reactome_2022” database. Pathways with an adjusted p-value < 0.05 were considered significant, according to the enrichR statistical method based on computing the deviation from the expected rank for terms.

For the analysis of mRNA enrichments, only mRNA genes measured in the TG, TCGA and GLASS cohorts were considered. Enrichments were calculated for all significant genes in TG, TCGA, and GLASS cohorts. Additional enrichments were run for all filtered miRNA target genes, lncRNA target genes and genes significantly differentially expressed in both TG and GLASS cohorts as well as in all three cohorts.

#### Gene Regulatory Network analysis

Functional interactions, derived from annotated interactions and interactions predicted by a naïve Bayes classifier[36] were downloaded from Reactome and filtered for genes showing significant differential expression in our cohort. All the ncRNA - mRNA interactions that were filtered using correlation were added to the interactions. To assign each mRNA to a Reactome pathway category, all individual pathways within each of the 29 Reactome categories were first aggregated. For each category, a list of unique genes involved in any of its individual pathways was compiled. An mRNA was then assigned to a pathway category if its corresponding gene was present in the gene set associated with that category. A mRNA can be part of several categories. For the scoring of the network genes, we used the formula (|log2FC| / max(|log2FC|)) * (Node Degree / max(Node Degree)). The networks were visualised with ggraph version 2.1 and Cytoscape version 3.10.1. For the visualisation of genes with several annotated Reactome categories, the following hierarchy was followed, meaning that the gene was included in the first annotated category: cell cycle or DNA repair, ECM, signal transduction, and metabolism. Cell cycle and DNA repair categories as well as metabolism and metabolism of proteins categories were combined. A neural-associated category was created by combining the neuronal system Reactome category with genes implicated in neural functions from the developmental biology, vesicle-mediated transport, signal transduction categories, and non-annotated genes.

### Analysis of TCGA data

#### TCGA miRNA-seq, RNA-seq and CNAs

Raw unstranded Illumina HiSeq hg38 gene expression counts and raw miRNA expression counts, miRBase 22 for the TCGA-GBM and TCGA-LGG projects were acquired from the GDC database through TCGAbiolinks R library v.2.28.2 [37–39]. According to the WHO 2021 classification of CNS tumors, IDHwt grade 2 and 3 tumors with chromosome 7 gain and chromosome 10 loss, TERT promoter mutation or EGFR amplification were considered as IDHwt GBs [30]. For tumors with more than one data entry, one of the samples was randomly selected. The cohort consisted of 577 TCGA primary glioma patients with gene expression data. For the miRNA-seq, the cohort consisted of a total of 429 TCGA primary LGG and GBM cases. The raw unstranded counts were combined, normalized and differential gene expression was calculated using DESeq2 v.1.40.1 (grade 2-3 vs. grade 4) for miRNAs and mRNAs separately. Genes with an adjusted p-value below 0.05 (BH method) and |log2foldchange| > 1 were considered significant. For differential gene expression analysis, only primary IDHmut astrocytoma cases were considered. For violin plots, all available primary diffuse glioma samples were included in the normalisation.

Copy number alterations were downloaded from cBioPortal [40,41] (merged LGG and GBM cohorts [42]) and the cases were filtered for primary cases and grouped based on WHO2021 classification (see above).

### Analysis of GLASS data

#### GLASS RNA-seq and CNAs

Transcript-level matrix as well as case information for all samples was downloaded from synapse.org. Only patients with matched RNA-seq data from IDHmut astrocytoma tumors before and after progression to grade 4 were included. Cases with no treatment before progression and one case having received only TMZ before progression were excluded, resulting in tumors from eight patients used for the analysis. All the patients have received at least radiation. Three cases have received only radiation, two have received radiation while chemotherapy is unknown and three cases have received radiation and TMZ treatment. The transcripts were summed up using ensembldb (Ensembl v. 75 according to GLASS paper) and the tximport R package version 1.28.0. DESeq v.1.40.1 was used for normalization and calculation of differential expression.

Copy number alteration data was downloaded from synapse.org and filtered for the cases included in the RNA-seq analysis.

#### RNA-seq data from Sojka et al. (GSE276176)

The “GSE276176_hCS_feature_counts.xlsx” file was downloaded from GSE276176 and the data was normalised using DESeq2 v.1.40.1.

#### Statistical analysis

Statistical testing was performed with R version 3.6.2 (DE analysis, BiocManager version 3.10 for DESeq2 org.Hs.eg.db) and R version 4.3.3 for the remaining analysis. The statistical tests are specified within the main text and figure legends. Unless stated otherwise, all tests were conducted using a two-tailed approach.

## Results

We utilized strand-specific RNA-seq and small RNA-seq data originating from the same tissue samples from six patients (TG01–TG06) (hereafter referred to as the TG cohort) to elucidate the role of ncRNAs in the progression of grade 2–3 IDHmut astrocytomas to grade 4 after treatment. The whole-genome sequencing data generated from TG01–TG05 (previously reported in [7]) was used for DNA copy number alteration (CNA) analysis. Due to only targeted DNA-seq being available, CNAs from TG06 were not analysed.

### Clinical courses and mutations detected in the cases

The clinical courses and detected genomic alterations have been described in detail by Rautajoki et al. [7]. Briefly, tumors TG01–TG06 represented astrocytomas harboring a missense hotspot mutation in *IDH1* at p.R132 and alterations in *ATRX. TP53* was inactivated via missense or truncating mutations in all cases except for TG03, both before and after progression. TG01 showed a hypermutator phenotype associated with full inactivation of the *MSH2* and *DNMT3A* genes. Upon progression, either *CDKN2A* or *RB1* was inactivated in all the cases. In three cases, alterations in growth factor receptors were observed concomitantly with *CDKN2A* inactivation. The time between the primary surgery and progression to grade 4 ranged from 20 to 146 months (Fig. S1c). All patients had died at the start of the study.

All grade 4 tumors within our cohort were originally diagnosed as secondary glioblastomas; however, the diagnosis was later updated to grade 4 astrocytomas based on the World Health Organization (WHO) 2021 guidelines. All patients received radiotherapy, and all except TG04 chemotherapy before progression to grade 4 (Fig. S1c).

The results were analyzed together with public IDHmut astrocytoma data generated by the GLASS (including longitudinal, matched samples in the treatment setting) and TCGA (including only primary tumors) consortia for validation and comparison purposes (Fig. 1a). The TCGA cohort allowed the detection of grade-related changes independent of treatment influence within a larger cohort, whereas the GLASS cohort served as a partial validation of post-treatment changes. In the GLASS cohort, the comparison was always primary vs. grade 4 tumors, whereas in the TG cohort, it was before vs. after tumor progression to grade 4. In both TG and GLASS cohorts, all the cases have received at least radiotherapy before tumor progression to grade 4, however, the chemotherapy treatment differs. In GLASS, a higher fraction of patients received radiotherapy only (3/8 cases [and unknown chemotherapy status for two cases] vs. 1/6 in the TG cohort) (Fig. 1a). In the TG, no patient received TMZ before progression, while in the GLASS at least three patients received TMZ before progression. While TMZ primarily generates point mutations [43,44], the treatments used in the TG cohort also crosslink DNA and lead to double-strand breaks [45].

**Fig. 1.**
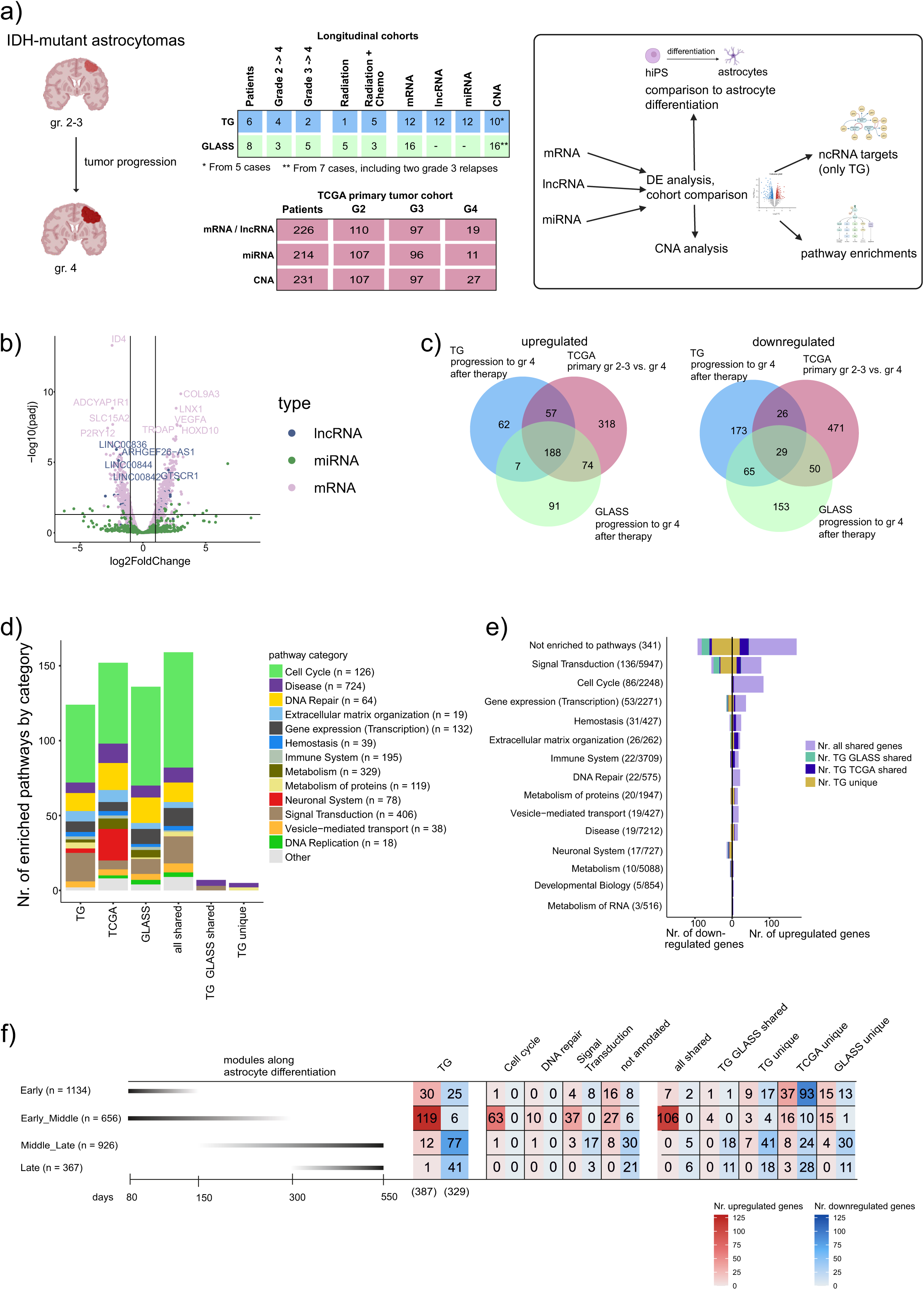
Differentially expressed protein coding genes are associated with increased cell proliferation, extracellular matrix organization, and decreased cell differentiation. **a)** Overview of the project and the used datasets. In-house longitudinal data from six patients (TG cohort) was analyzed in the context of public data from TCGA (representing only primary tumors) and GLASS (only for mRNA comparison). The GLASS cohort includes longitudinal data from eight IDHmut astrocytoma cases that have progressed to grade 4 after therapy, but the received treatments partly differed between TG and GLASS. Differentially expressed genes were also compared to astrocyte differentiation-related genes from [50]. **b)** Altogether 398 upregulated and 331 downregulated mRNAs, lncRNAs, and miRNAs were differentially expressed upon post-therapy progression in TG. **c)** Comparison of differentially expressed mRNAs in TG, TCGA, and GLASS datasets reveals that a higher fraction of upregulated (188, 59.9%) than downregulated (29, 9.9%) genes are shared between all three cohorts. The TCGA cohort includes only primary tumors and was used as a reference to separate post-treatment-related changes from global grade-related differences in IDHmut astrocytomas. Seven upregulated and 65 downregulated genes were detected in both the TG and GLASS cohorts, so recurrently in the post-treatment setting. All the cases in the TG and GLASS cohorts received radiotherapy before progression. **d)** When running the pathway enrichment analysis with DE mRNAs measured in all three cohorts, the cell cycle pathway category includes the highest number of enriched pathways, followed by DNA-repair and signal transduction categories. Each pathway significantly enriched by respective DE genes was assigned a pathway category based on the Reactome Pathway Hierarchy. Categories with less than three enriched pathways across all comparisons are categorized as “Other”. N-numbers in brackets indicate how many pathways in total belong to each category based on Reactome annotations. **e)** The highest number of DE genes in the TG cohort fall within enriched pathways in signal transduction, cell cycle, and ‘gene expression (transcription)’ categories. Nearly all of the genes in Cell cycle and DNA repair categories were DE also in TCGA primary tumor cohort and GLASS. Half (56%) of DE mRNA genes are not part of any enriched pathway (marked as ‘not annotated’)). Numbers in brackets indicate the number of DE mRNAs and the total number of unique genes mapped to each category based on Reactome. **f)** The Early_Middle WGCNA module as reported by Sojka et al. [50] includes the highest fraction of TG DE genes, which are mostly upregulated, shared among all three datasets, and contain a high fraction of cell-cycle related genes. Within the Middle_Late and Late modules, the majority of overlapping genes are downregulated and unique to the post- treatment setting.

### Transcriptome changes upon progression to grade 4 are dominated by cell cycle and signal transduction

The TG cohort (Fig. S1c,d) served as a discovery cohort, in which the transcriptomic changes between matched grades 2–3 and post-treatment grade 4 IDHmut astrocytomas were defined using paired DESeq2. This revealed 676 differentially expressed (DE) mRNAs (adj. p < 0.05, |log2FC| > 1), 33 lncRNAs (adj. p < 0.05, |log2FC| > 1), and 20 miRNAs (adj. p < 0.05, |log2FC| > 1), of which 398 were upregulated and 331 downregulated (Fig. 1b).

To validate our findings, we explored the landscape of DE transcripts detected in the GLASS longitudinal and TCGA primary tumor cohorts. Because only a few ncRNAs were quantified in the GLASS cohort, we restricted this analysis to protein-coding genes (mRNAs) measured in all three cohorts, covering 314 upregulated and 293 downregulated mRNAs in TG.

Our similar DE gene analysis between grades 2–3 and grade 4 IDHmut astrocytomas in the TCGA and GLASS datasets (Fig. S1e, Table S1-2) uncovered 217 genes (188 upregulated and 29 downregulated, 35.7%) demonstrating similar significant changes in all three cohorts (Fig. 1c, Table S3). Altogether, 72 genes (7 upregulated and 65 downregulated, 23.5%) were detected in the post-treatment setting (DE in both the TG and GLASS cohorts) (Fig. 1c, Table S2). A high fraction (245/314, 78%) of mRNAs upregulated in TG was also upregulated in grade 4 primary tumors in TCGA, whereas downregulated genes were mainly either TG unique (173/293, 59%) or detected after therapy in both TG and GLASS (65/293, 22%) (Fig. 1c).

Next, we calculated significantly enriched Reactome pathways (adj. p < 0.05) based on DE genes (Tables S4, S5 and S6) and divided them into categories based on the Reactome Pathway Hierarchy. Most of the enriched pathways belonged to the cell cycle category, followed by DNA repair and signal transduction (Fig. 1d, Tables S7 and S8). Many DNA repair genes are upregulated during mitosis and show a strong correlation with cell cycle-related genes [7,46]; therefore, their enrichment is likely related to increased proliferation. The upregulation of proliferation was further indicated by the upregulation of 13 previously reported G1/S-phase and 34 G2/M-phase markers (Table S3) [47].

Next, we investigated which DE mRNAs in our TG cohort contributed to the observed enrichment by combining the genes from each pathway category and calculating their overlap with DE genes. Extracellular matrix (ECM) organization showed the highest fraction of DE genes compared to the total number of unique genes in that category (26/262, 9.92%) (Fig. 1e). Most genes in the cell cycle and DNA repair categories were upregulated across the three cohorts, suggesting that they represented treatment-independent tumor grade-related processes (Fig. 1e). A majority of DE mRNAs were not part of any enriched pathway (341/607, 56.2%) and included a high fraction of the downregulated genes (200 of 293 downregulated genes, 68.3%) (Fig. 1e).

Overall, these results show robust upregulation of proliferation and mitosis-associated DNA repair with increasing tumor grade. These results are consistent with those of previous reports, in which cell cycle and mitosis were significantly enriched terms for upregulated genes in IDH-mutant glioma progression [7,8]. Furthermore, the downregulated genes were mostly associated with post-treatment tumor progression.

### Genes upregulated in astrocyte differentiation are mostly downregulated upon tumor progression

Dedifferentiation is a process that occurs upon tumor progression [48,49]. Therefore, we explored the number of our DE genes linked to astrocyte differentiation. Sojka et al. differentiated human induced pluripotent stem cells within cortical organoids into astrocytes over 550 days, while collecting regular RNA-seq measurements [50]. We overlapped our DE genes with the genes included in the weighted correlation network analysis modules, each representing genes with correlating RNA-based expression changes along the differentiation process (Fig. 1f, Fig. S1f). In total, 352 DE genes (49.2% of all DE genes) in the TG cohort were included in the modules, with each gene assigned to only one module. Altogether, 119 of genes included in the Early_Middle module (highest expression until day 300 [50]) were upregulated upon progression in our TG data, and 106 (89%) of them show upregulation also in the TCGA and GLASS datasets, including 62 cell cycle-related genes based on the Reactome annotations (Fig 1f, Table S9). Meanwhile, the majority of TG DE genes in the Middle_Late (upregulated after day 150) and Late (upregulated after day 350) modules were downregulated upon progression (77/89 [86.5%] and 41/42 [97.6%], respectively). Interestingly, their downregulation was detected especially in the post-treatment setting (59 genes unique to TG and 29 genes shared with GLASS; 50% and 24.6% of downregulated genes within these two modules, respectively). Middle_Late and Late modules included 59 genes that were not annotated in Reactome, 51 (86.4%) of which were downregulated upon tumor progression, suggesting that their downregulation was associated with lowered astrocytic cell differentiation. The lower differentiation state was further supported by the upregulation of immature astrocyte markers, e.g. *NES* [51] and *THBS1* [52] uniquely in the TG cohort.

Overall, grade 4 tumors showed a signature of cell dedifferentiation upon progression, especially in the post-treatment setting.

### Dysregulated *NEAT1*, *PVT1*, and *CYTOR* lncRNAs are predicted to regulate genes implicated in cell proliferation, nervous system development, and ECM organization

Because lncRNAs regulate different key processes in health and disease [53], we investigated which lncRNAs were dysregulated during tumor progression and to what extent they contributed to the observed transcriptional phenotypes. In the TG cohort, 19 upregulated and 14 downregulated lncRNAs were detected after progression to grade 4 (Fig. 2a, Table S10). Most of the observed DE lncRNAs were unique to our cohort, as only seven upregulated and four downregulated lncRNAs were also DE between primary grades 4 and 2–3 IDHmut astrocytomas in the TCGA tumor cohort (Fig. 2b). These shared genes included upregulated *CYTOR* and *PVT1*. Among the 33 DE lncRNAs, only the expression of COLCA1 was measured in the GLASS cohort (and was similarly downregulated: LFC -2.1, adj. p = 0.005); therefore, only CNA data from the GLASS cohort was used in the lncRNA analysis.

**Fig 2.**
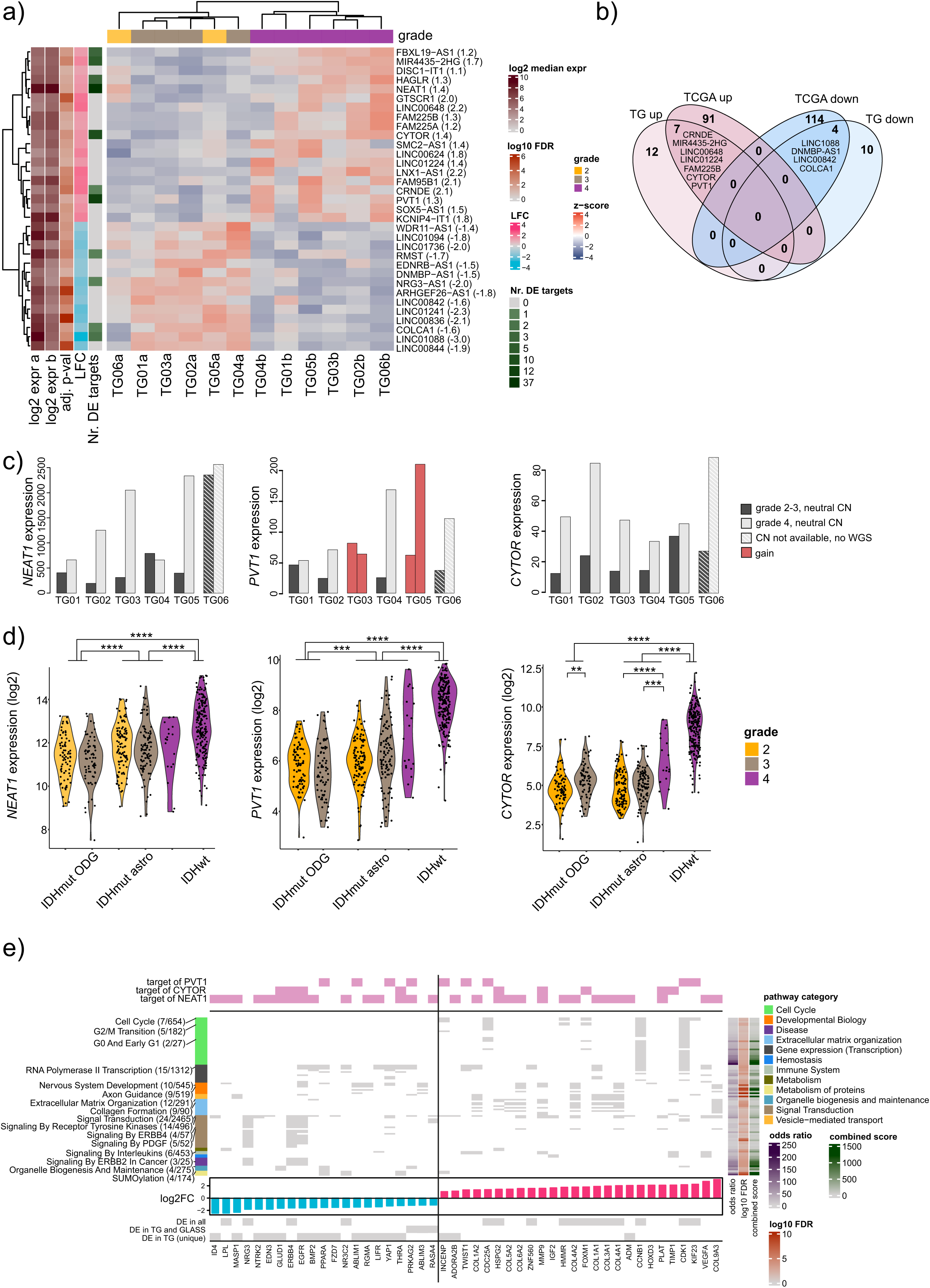
Predicted targets of tumor progression-associated lncRNAs are involved in cell cycle, signal transduction, and extracellular matrix organization. **a)** The detected 19 upregulated and 14 downregulated lncRNAs show significant expression changes upon tumor progression. Log2 median expression shows the normalized log2 transformed counts of each lncRNA in both lower grade and progressed tumor. Log10 false discovery rate (FDR) indicates the adj. p-value of the differential expression, transformed by log10. LFC refers to the log2FC of the lncRNA detected after progression to grade 4. “Nr. DE targets” indicates the amount of correlating DE mRNA targets observed for each lncRNA. Grade refers to the World Health Organization tumor grade of the sample. The z-score represents gene expression values scaled across samples. **b)** Comparison against the DE lncRNAs in TCGA primary tumor dataset reveals seven commonly upregulated lncRNAs, including *PVT1* and *CYTOR,* as well as four shared downregulated lncRNAs. **c)** lncRNAs *NEAT1*, *CYTOR*, and *PVT1* are upregulated upon progression in the TG cohort. One-copy gain of *PVT1* is detected in TG03 and TG05 both before and after progression. Expression refers to normalized counts. d) *NEAT1*, *PVT1*, and *CYTOR* show the highest expression in IDHwt GBs and lowest in IDHmut oligodendrogliomas (IDHmut ODG) in TCGA. *NEAT1* expression does not differ significantly between IDHmut astrocytoma grades. Expression (log2) refers to log2 normalized counts. ** p < 0.01, *** p < 0.001, **** p < 0.0001, adj. Wilcoxon rank-sum test. **e)** The predicted DE targets of the progression-related lncRNAs are associated with the cell cycle, developmental biology, extracellular matrix organization, and ERBB signaling. A majority of the genes (85.1%) are predicted to be regulated by one of the three highest-scoring lncRNAs (*NEAT1*, *CYTOR*, or *PVT1*). Pathway category indicates to which Reactome pathway category each pathway belongs. Log10 FDR indicates the log10 transformed adj. p-value of the enrichment. The combined score is derived as the product of the odds ratio and the log10 FDR. The target genes’ LFC values in the TG cohort as well as DE status in different cohorts are visualized at the bottom.

The DE lncRNAs in the TG cohort were linked to their targets based on the LncTarD and NcPath databases. Target genes needed to be DE (adj. p < 0.05) (Fig. 2a, Table S10) and show either a negative or positive correlation (|Spearman’s and Pearson’s correlation coefficient| > 0.5) with the upstream lncRNA. All target genes showed clear differences in expression (|log2FC| > 0.88). In total, the 33 lncRNAs were predicted to regulate 62 DE genes (0–38 targets per lncRNA) (Fig. S2a).

DE lncRNAs were scored based on their absolute log2FC and the number of predicted DE targets (see Methods). The lncRNA with the highest score was *NEAT1* (log2FC 1.4) (Fig. 2c). *NEAT1* is overexpressed in glioma tissue, and a high expression has been associated with larger tumor size and increased risk of tumor recurrence. High *NEAT1* expression correlated with poor OS in high-grade (grades 3–4) gliomas, as well as in cases that received postoperative chemotherapy.[54] *NEAT1* has also been linked to proliferation and chemoresistance [55]. The next highest score was for *PVT1* (log2FC 1.3) (Fig. 2c). In addition to proliferation, it has been implicated in chemotherapy and radiation resistance across cancers [56]. *CYTOR* (log2FC 1.4) is upregulated across cancers, correlates with shorter OS in various cancers, and is implicated in tumorigenesis and proliferation [57,58]. It had the third-highest score in our dataset. All three highest-scoring lncRNAs were upregulated after disease progression (Fig. 2c). Among IDHmut astrocytomas, grade 4 primary tumors showed the highest *CYTOR* expression (Fig. 2d). *PVT1* was statistically significant only when grade 2 and grade 3 tumors were combined and compared to grade 4 tumors in DESeq2 using the Wald test (Fig. 2b and Fig. 2d). Interestingly, *NEAT1* was significantly upregulated in grade 4 IDHmut astrocytomas only in the post-treatment setting (Fig. 2d).

Next, we investigated whether DNA copy number alterations influenced the observed changes in expression. Among the three highest-scoring lncRNAs, only *PVT1* gains (on chr8 q24.21) were significantly associated with increased expression in IDHmut astrocytomas and IDHwt GBs in the TCGA cohort (Table S11, Fig. S2e). In our cohort, *PVT1* is gained in two cases, but the gains are not progression-associated (Fig. 2c). According to GLASS data, *PVT1* showed two progression-associated gains (Table S12).

Finally, we performed gene set enrichment analysis for the 62 predicted DE target genes, resulting in 47 genes that were significantly enriched in 97 pathways from the Reactome database (Fig. 2e, Table S13). The majority of DE genes (40/47, 85.1%) were downstream of *NEAT1*, *CYTOR*, and/or *PVT1* (Fig. 2e). *NEAT1* was predicted to regulate the highest number of genes (n=38), including eight collagen subunits, contributing to the enrichment of ECM organization (Fig. 2e). Experimentally validated *NEAT1* target *COL1A1* [59] (log2FC = 1.9, Pearson’s corr. = 0.66, Spearman’s corr. = 0.78) is required for the formation of mesenchymal-like multicellular fascicles of elongated and aligned cells (called oncostreams), which are important for glioma invasion and spread. Both *COL1A1* and *MMP9* (log2FC 1.6, Pearson’s corr. = 0.69, Spearman’s corr. = 0.72, experimentally validated in [60]) are overexpressed in the oncostreams [61]. The presence of oncostreams and higher *COL1A1* expression is associated with a higher grade of diffuse glioma [61].

Consistent with *NEAT1* being upregulated only in the post-treatment setting, 17 of the 38 (45%) *NEAT1* target mRNAs were DE only in the TG cohort or shared between TG and GLASS. All but two (*FOXM1* and *TIMP1*) of 10 predicted *CYTOR* targets were shared with *NEAT1*, indicating a potential cooperative involvement especially in signal transduction. *PVT1* targets were distinct and have been implicated in the cell cycle, including the experimentally validated targets *KIF23* [62] (log2FC 2.3, Pearson’s corr. = 0.87, Spearman’s corr. = 0.6) and *CDK1* [63,64] (log2FC = 2.2, Pearson’s corr. = 0.92, Spearman’s corr. = 0.6). *PVT1* has been reported to regulate *CDK1* through various miRNA interaction partners, which vary depending on the cancer type [63,64].

### Involvement of DE miRNAs in cell cycle and signal transduction

Small RNA-seq data was generated from all samples in the TG cohort and utilized for miRNA analysis (Table S14). To identify the putative targets of the 12 upregulated and 8 downregulated miRNAs, we obtained targets from miRWalk and mirTarBase. Target genes needed to be DE (adj. p < 0.05) and show a negative correlation (Spearman’s and Pearson’s correlation coefficient < −0.5) with the upstream miRNA. This resulted in a total of 70 DE genes (0–18 per miRNA) predicted to be regulated by at least one miRNA (Fig. 3a, Fig. S3a, Table S15).

**Fig 3.**
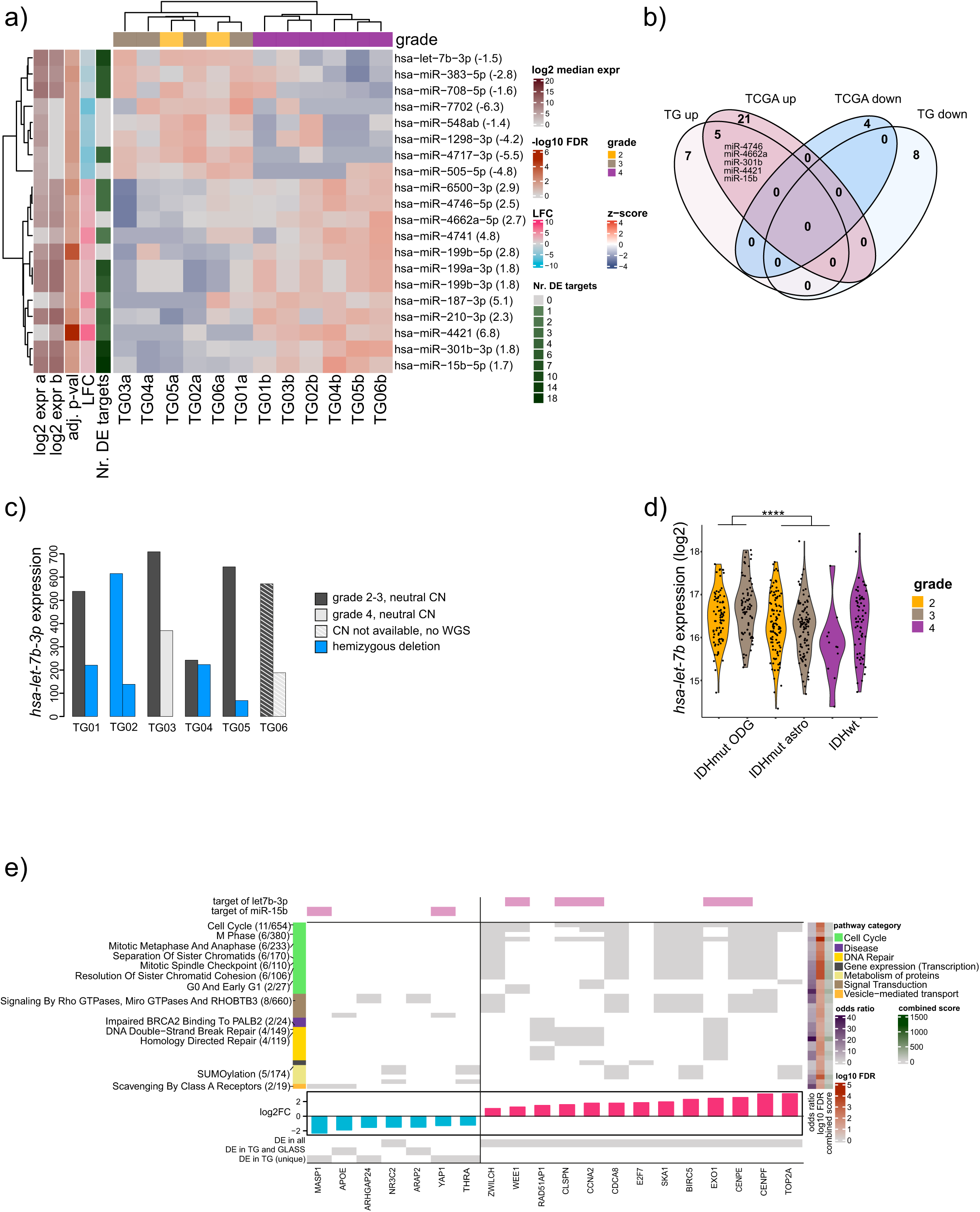
Predicted targets of tumor progression-associated miRNAs are involved in cell cycle, signal transduction, and DNA repair. **a)** Upon tumor progression, 20 miRNAs were dysregulated, out of which 12 were upregulated and 8 were downregulated. Log2 median expression shows the normalized log2 transformed counts of the miRNA in both lower grade and progressed tumor. Log10 false discovery rate (FDR) indicates the adj. p-value of the miRNA, transformed by log10. LFC refers to the log2FC of the miRNA detected after progression to grade 4. “Nr. DE targets” indicate the amount of correlating DE mRNA targets observed for each miRNA. Grade refers to the World Health Organization tumor grade of the sample. The z-score represents gene expression values scaled across samples. **b)** Five of the upregulated miRNAs are also upregulated in primary grade 4 vs. grade 2–3 IDHmut astrocytomas in the TCGA dataset. None of the downregulated miRNAs are shared with TCGA. **c)** The miRNA *hsa-let-7b-3p* shows recurrent one-copy deletions and downregulation upon tumor progression in the TG cohort. Expression refers to normalized counts. **d)** The expression of *hsa-let-7b* in TCGA primary tumor cohort. No significant differences in expression are detected between different tumor grades representing the same tumor type. Expression (log2) refers to log2 normalized counts. *p < 0.05, ***p < 0.001, ****p < 0.0001, adj. Wilcoxon rank-sum test. **e)** Differentially expressed miRNA targets are enriched for cell cycle and signaling Reactome pathways. Five of 11 cell cycle and DNA repair-related genes are targeted by *hsa-let-7b-3p*. Log10 FDR indicates the log10 transformed adj. p-value of the pathway. The combined score is derived as the product of the odds ratio and the log10 FDR. The target genes’ log2FC values in the TG cohort as well as DE status in different cohorts are visualized at the bottom.

Five of the upregulated miRNAs, including *hsa-miR-15b* showed similar expression patterns in the TG and TCGA primary tumor cohorts (Fig. 3b) (Table S16). None of the downregulated miRNAs were shared between the TCGA and TG analysis. miRNA sequencing was not performed in the GLASS cohort; therefore, no comparisons were made. Based on DNA CNA analysis in the TG cohort, recurrent progression-related hemizygous deletions coupled with downregulation were detected in *hsa-let-7b-3p* (on chr22 q13.31, log2FC = −1.5) (Fig. 3c), which is involved in cell cycle regulation [65]. *hsa-let-7b-3p* was lost in three cases after progression (Fig. 3c) as well as in both tumors from TG02 (Fig. 3c). In the TCGA primary tumor cohort, *hsa-let-7b* did not show differential expression between IDHmut astrocytoma grades (log2FC = −0.4, adj. p = 0.38) (Fig. 3d), linking the downregulation to the matched setting. However, it experienced a recurrent DNA copy number loss which was also associated with lower expression in primary IDHmut astrocytomas (Fig. S3b). The frequency of gene loss in TCGA is increased with tumor grade:

*hsa-let-7b-3p* was homozygously deleted in four patients, and hemizygous deletions were observed in 13/12/48% of grade 2/3/4 IDHmut astrocytomas, respectively (Table S17). In addition, within the GLASS cohort, a hemizygous deletion was observed in two cases upon progression (Table S18).

The miRNA *hsa-miR-15b-5p* had the highest score based on its absolute log2FC and the number of predicted, correlating DE targets (log2FC = 1.7, 18 targets). Its high expression is associated with worse survival in gliomas [66,67] and its inhibition has induced apoptosis in a GB cell line [68]. Fittingly, it is upregulated in both the TG and TCGA IDHmut astrocytoma cohorts (Fig. S3c, Fig. S3d). *hsa-miR-15b-5p* (in chr 3q25.33) had a one-copy gain in one progressed TG case, and its one-copy gain was associated with higher expression in IDHmut astrocytomas in the TCGA cohort, based on the three cases for which expression data was available (Fig. S3e).

Considering the 35 significantly enriched Reactome pathways for the predicted DE miRNA target genes (Fig. 3e, Table S19), 22 of the 35 (62.9%) pathways belong to the cell cycle or DNA repair categories. *hsa-let-7b-3p* was predicted to regulate 5 of 12 (42%) target genes in the enriched pathways, such as the experimentally validated *CCNA2* [69] (log2FC = 1.8, Pearson’s corr = −0.64, Spearman’s corr. = −0.71) (Fig. 3e).

In conclusion, recurrent CNA losses are associated with *hsa-let-7b-3p* downregulation, which was significant particularly in the post-treatment setting.

### Both miRNAs and lncRNAs were predicted to contribute to the increased cell proliferation and downregulated neural functions

To investigate the involvement of lncRNAs and miRNA targets in the same cellular processes, we reconstructed a GRN based on the Reactome interactions of TG DE genes and filtered them for DE ncRNAs and their direct DE interaction partners (Fig. 4a, Fig. S4a, Table S20). A total of 34 mRNAs were targeted by more than one ncRNA, and six of these were targeted by at least one lncRNA and one miRNA. Most miRNA target genes were implicated in only one process; for example, *hsa-miR-708-5p* and *hsa-miR-4717-3p* targets were all in the cell cycle category (Fig. 4a-b, Fig. S4a). By contrast, lncRNA target genes may be more pleiotropic in nature; for example, *CYTOR*, *PVT1*, and *NEAT1* targets belong to at least five different Reactome categories. *NEAT1* interacted with 13 of 14 upregulated mRNA target genes in the ECM category (Fig. 4c), linking it to altered ECM in grade 4 IDH-mutant tumors. While several miRNAs and lncRNAs are predicted to regulate genes related to cell cycle (Fig. 4b) and signal transduction (Fig. S4b), in both of these categories, none of the mRNAs are targeted by both miRNAs and lncRNAs simultaneously.

**Fig 4.**
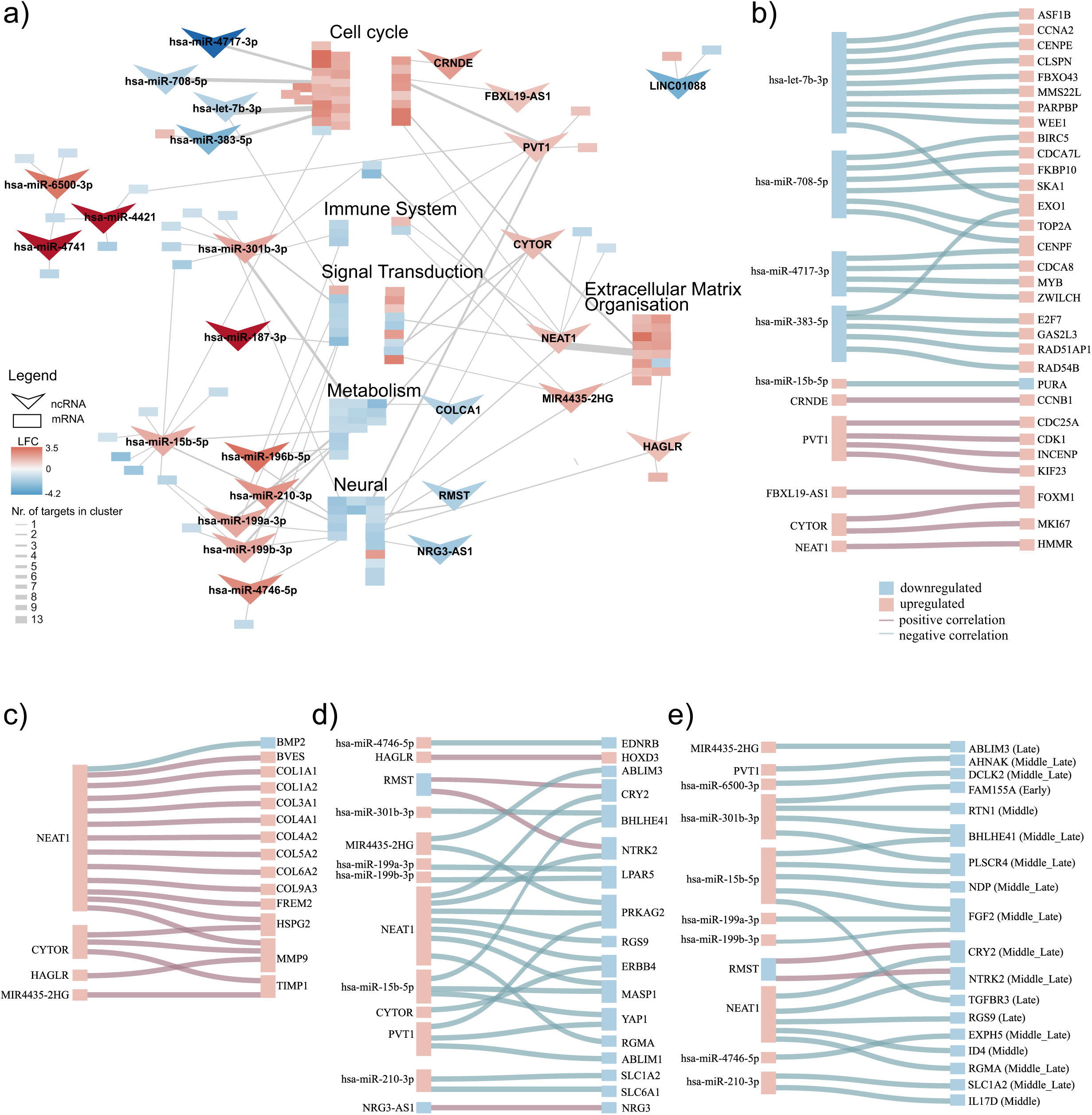
Both miRNAs and lncRNAs target genes in major pathway categories. **a)** A simplified gene regulatory network based on Supplementary Figure 4, consisting of 125 DE mRNAs predicted to be regulated by 25 ncRNAs. The color of the node indicates the log2FC. A V shape indicates an ncRNA and a rectangle indicates an mRNA. Reactome annotations of clusters are marked above them. Connections between ncRNAs and the clusters are marked if at least one predicted target mRNA belongs to the respective cluster. **b)** 31 DE mRNAs involved in cell cycle and DNA repair are regulated by five miRNAs and five lncRNAs. None of the targets are shared between miRNAs and lncRNAs. ncRNAs are on the left side of the figure, the predicted targets are on the right side. The colour of the line connecting them indicates positive or negative correlation. The colour of the rectangle indicates a positive or negative log2FC post-progression in the TG cohort and the size of the ncRNA box indicates the amount of predicted targets. **c)** Four lncRNAs are predicted to target 14 DE mRNAs involved in ECM organisation. *NEAT1* is predicted to regulate 13 of the 14 targets. **d)** Among the DE neural genes, two are predicted to be targeted by miRNAs and lncRNAs simultaneously. An additional six targets are putatively regulated by five miRNAs and ten other targets by five lncRNAs. **e)** Altogether, 18 neural or non-annotated ncRNA targets (all downregulated) are part of one of the five WGCNA RNA modules describing astrocyte differentiation as reported by Sojka et al. The module of the respective mRNA is indicated in brackets behind its name.

Within the GRN, two categories stood out, with the majority of genes being downregulated upon progression: metabolism (Fig. S4c) and neural genes (Fig. 4d). To analyze the involvement of ncRNAs in neural activity in the progression setting, we created a neural-associated category (Table S21, see Methods). Seven lncRNAs (including *NEAT1* and *PVT1)* and six miRNAs (including *hsa-miR-15b-5p*) were predicted to regulate at least one of the 17 neural-associated targets (Fig. 4d). With the exception of *HOXD3* (log2FC 2.1), all target genes were downregulated. To determine the overlap with astrocyte differentiation-related genes, we extracted mRNAs within our neural-associated category as well as non-annotated mRNAs that are part of one of the differentiation modules by Sojka et al., revealing 18 downregulated ncRNA targets, of which 14 were part of the Middle_Late or Late module (Fig. 4e). Seven of the 18 targets were shared with our neural category, including six annotated by us, confirming their relevance to neural processes.

In addition to the previously mentioned Reactome category-based mRNA clusters, an interesting subcluster consisted of upregulated *hsa-miR-4421* (log2FC 6.8), *hsa-miR-6500- 3p* (log2FC 2.9), and *hsa-miR-4741* (log2FC 4.8,), which were predicted to target the downregulated gene *NT5DC3* (log2FC = -1.2, Pearson’s corr. = −0.65/−0.67/−0.62, Spearman’s corr = −0.82/−0.87/−0.66, respectively, predicted) (Fig. S4a). Most of the genes within the above-mentioned subcluster are neither annotated within Reactome nor well studied in the literature. *hsa-miR-6500-3p* was predicted to also regulate the downregulated *TDRD3* (log2FC = −0.99, Pearson’s corr = −0.59, Spearman’s corr = −0.64, predicted) and *DCLK2* (log2FC −0.9, Pearson’s corr = −0.64, Spearman’s corr = −0.67, predicted) genes.

Although the role of *NT5DC3* is largely unknown, both *TDRD3* and *DCLK2* were involved in neural processes, such as synaptic plasticity and neuronal injury response [70,71], indicating that this subcluster might also play a role in neural activity.

To summarise, both miRNAs and lncRNAs are associated with the upregulation of cell-cycle related genes and the downregulation of neural function-related genes that are linked to cell dedifferentiation.

## Discussion

This study describes the phenotypic changes that occur in IDHmut astrocytomas upon progression after treatment as well as the dysregulated ncRNAs involved in this process. To the best of our knowledge, no ncRNA sequencing data from matched IDHmut astrocytoma cases before and after progression has been published.

One underexplored aspect of ncRNA research in the context of tumor progression is the influence of copy number alterations. Herein, we report the downregulation of *hsa-let-7b-3p* after post-treatment progression coupled with hemizygous losses. Recurrent progression-related losses were detected in both TG and GLASS cohorts, and they were also more frequent in primary grade 4 than 2-3 tumors in TCGA data. An association between let-7b deletion and lower mature let-7b expression has also been reported in ovarian cancer [72]. Notably, significant grade-related changes in *hsa-let-7b-3p* expression were observed only in the matched post-treatment setting. Interestingly, *hsa-let-7b* is downregulated after radiation treatment in fibroblasts [73]. *hsa-let-7b* is also downregulated in glioma relapses, even if the tumor grade remains stable, with the lowest expression observed in high-grade tumors [74]. Mechanistically, *hsa-let-7b-3p* is predicted to suppress the expression of genes involved in cell proliferation, such as the upregulated *CENPE*, *CCNA2, WEE1, EXO1*, and *CLSPN* genes, consistent with its role as a tumor suppressor and inhibitor of cell proliferation [75]. CCNA2 plays a crucial role in DNA synthesis during the S phase by binding to CDK2 [76] and activating CDK1 during the G2-phase, influencing chromatin condensation [77]. *CCNA2* has been experimentally validated as a *let-7b target* in gliomas [69]. Although our *let-7b* targets did not fall within the neural category, let-7 promotes astrocytic cell differentiation in both normal and glioma cells [78,79]. The relevance of let-7b is further highlighted by reports on its prognostic role in gliomas [80]. Treatment of patient-derived glioma stem cells with phenformin increases let-7 expression, leading to decreased glioma stem cell self-renewal [81]. Overall, *hsa-let-7b-3p* has emerged as an interesting target for further studies on gliomas in the context of treatment resistance and post-treatment progression.

An increased proliferation index is a characteristic of diffuse astrocytoma progression and is also previously detected in the TG cohort together with recurrent inactivation of cell cycle checkpoints after progression [7,82]. Therefore, it is not surprising that, in addition to *hsa-let- 7b-3p*, several other DE ncRNAs have been implicated in cell cycle regulation. We observed upregulation of *PVT1*, which promotes cell proliferation [83] and TMZ resistance in gliomas [84]. Based on the previous experimental data, *PVT1* targets *CDK1* [63,64], which has a crucial role in the cell cycle by promoting the G2/M and G1/S transitions, as well as G1 progression [85]. Its role is emphasized by it being the only essential cyclin dependent kinase in mammals [86]. In addition, the previously reported *PVT1* targets *INCENP* [87], and *KIF23* [62] were upregulated also in our analysis. Another upregulated key player in the cell cycle, *FOXM1*, is targeted by *CYTOR* and *FBXL19-AS1*, both of which have been reported in the context of proliferation before [88–90].

Polycomb repressive complex 2 (PRC2) subunit EZH2 writes the H3K27me3 modification, which is contributing to the malignancy of various cancers [91]. In gliomas, EZH2 has been observed to be overexpressed and its high expression predicts short OS [92]. Its inhibition increases the chemosensitivity [93,94] and induces cell cycle arrest [92]. Several DE lncRNAs have been reported to interact with EZH2. Although most of these interactions occur at the protein level and therefore are not detected with our approach, at least 5 of 33 DE lncRNAs (15%) interact with EZH2, namely *NEAT1* [95], *CYTOR* [96], *MIR4435-2HG* [97], *CRNDE* [98], and *PVT1* [99]. Among the DE miRNAs, 2 of 20 (10%) interact with EZH2, namely *hsa-let-7b* [100], and *hsa-miR-708* [101].

Our study cohort was limited in size and void of identical treatment schemes, partially because of the rarity of these tumors and the difficulty in accessing matching longitudinal samples. It is also challenging to generate IDHmut astrocytoma cell lines and use the few available ones for validation purposes. This all makes it more difficult to estimate result generalizations. However, matching increases the power of the analysis and provides information on tumor evolution over time. Furthermore, we extended our perspective and increased the sample size by analyzing publicly available data from TCGA and GLASS. TCGA allowed us to detect grade-related differences that were not treatment-dependent, whereas GLASS provided longitudinal mRNA data for comparison purposes.

While we selected strict criteria for ncRNA targets and highlighted especially experimentally validated targets, many of the predicted targets were not experimentally validated, and we cannot rule out the possibility that their upregulation or downregulation is also driven by other factors. To overcome this challenge, we, for example, focused on experimentally-validated proliferation-related targets that have a known role in cell cycle regulation, as mitosis is known to induce changes in the expression of numerous genes despite ncRNA-mediated regulation. Further studies are required to validate the computationally predicted ncRNA targets in the context of IDHmut astrocytoma progression.

This study identified dysregulated ncRNAs associated with post-treatment progression of IDH-mutant astrocytoma and explored their potential regulatory roles by characterizing their targets within a GRN framework. Many of the targets were involved in cell proliferation and neural functions, with implications to progression-related dedifferentiation. In addition to their differential expression, *PVT1* and *hsa-let-7b-3p* were affected by recurrent DNA CNAs. A subset of ncRNAs, such as *hsa-let-7b-3p* and *NEAT1*, has been previously reported in the context of treatment response and resistance in other cancers. Based on our results, ncRNAs indeed have a role in these clinically relevant processes.

**SFig 1. a)** The obtained RNA-seq data has good quality. Histograms include the frequency of each observed read count bracket, scatter plots indicate the expression values across genes. Pearson’s correlation values are calculated between sample pairs and are shown above the diagonal. **b)** Similar data quality plots generated from the miRNA sequencing data. **c)** Timelines of all patients in the TG cohort, including treatment information. All cases received radiation treatment before progression to grade 4, and all but TG04 adjuvant chemotherapy. Reprint with permission from [7]. **d)** A principal component analysis plot for the 12 sequenced samples reveals heterogeneity among the patients. **e)** Scatterplot showing the log2FC values of each gene that has an adj. p < 0.05 significant in at least one cohort in TG vs. GLASS and TG vs. TCGA comparison, respectively. **f)** Expression of TG DE genes that are included in WGCNA modules from Sojka et al. [50] along the astrocyte cell differentiation trajectory. The log2FC values in the TG cohort are visualized on the left side of each heatmap, revealing prevalent upregulation of Early_Middle module genes and downregulation of Middle_Late and Late module genes upon tumor progression.

**SFig 2. a)** Expression patterns of 62 DE target mRNAs that are predicted to be targeted by 11 DE lncRNAs in the TG cohort. Correlation indicates whether the correlation coefficient between lncRNA and mRNA is positive or negative. LFC indicates the log2FC of the respective mRNA and grade indicates the tumor grade. **b)** *PVT1* expression is associated with its DNA CNA state in TCGA data (0 refers to normal diploid copy number). *p < 0.05, **p < 0.01, ***p < 0.001, ****p < 0.0001, adj. Wilcoxon rank-sum test.

**SFig 3. a)** Expression patterns of 70 DE target mRNAs that are predicted to be targeted by 14 DE miRNAs in the TG cohort. Correlation indicates whether there is a negative correlation coefficient between miRNA and mRNA. Correlation indicates whether a significant negative correlation between mRNA and miRNA was observed. LFC indicates the log2FC of the respective mRNA, grade indicates the grade of the tumor. **b)** For *hsa-let-7b*, hemizygous losses are associated with lower expression in the TCGA cohort. 0 refers to normal diploid copy number. * p < 0.05, ** p < 0.01, adj. Wilcoxon rank-sum test. **c)** Upregulated miRNAs *hsa-miR-15b-5p* shows the highest scores based on the absolute log2FC and number of predicted targets. *hsa-miR-15b-5p* (in chr3q25.33) shows a one-copy gain in TG03b. Expression refers to normalized counts. **d)** *Hsa-miR-15b,* has the highest expression in grade 4 IDHmut astrocytoma tumors compared to grades 2 and 3 tumors or other tumor types. Expression refers to log2 normalized counts. *p < 0.05, **p < 0.01, ***p < 0.001, ****p < 0.0001, adj. Wilcoxon rank-sum test. **e)** One copy gain of *hsa-miR-15b* is associated with higher expression in IDHmut astrocytomas. 0 refers to normal diploid copy number. *p < 0.05, **p < 0.01, adj. Wilcoxon rank-sum test. Expression refers to log2 normalized counts.

**SFig 4. a)** A gene regulatory network consisting of 125 DE genes predicted to be regulated by ncRNAs. The color of the node indicates the log2FC. A V shape indicates an ncRNA and a rectangle indicates an mRNA. MiRNAs were ordered to the left side, whereas lncRNAs were at the right. For genes with several annotated Reactome categories, the following hierarchy was followed, indicating that the gene was included in the first annotated category as follows: cell cycle or DNA repair, extracellular matrix organization, signal transduction, and metabolism. The DNA repair category was fused to the cell cycle category and the metabolism of proteins category was combined with the metabolism category. Unannotated target genes are positioned outside of midline. We combined the neuronal system Reactome category with genes implicated in neural functions from the developmental biology, vesicle-mediated transport, signal transduction categories, and non-annotated genes. **b)** Six miRNAs target six mRNAs coding for genes involved in signal transduction. Four lncRNAs target an additional nine mRNAs. **c)** Five lncRNAs and five miRNAs regulate 14 mRNAs involved in metabolism. Three of the targets are shared between lncRNAs and miRNAs.

## Supplementary Tables

Table S1. Genes that are differentially expressed in grade 4 primary tumors vs. primary grade 2-3 tumors in the TCGA cohort.

Table S2. Genes that are differentially expressed in matched post-treatment grade 4 tumors vs. primary grade 2-3 tumors in the GLASS cohort.

Table S3. Genes that are differentially expressed in the TG cohort, with expression statistics in TCGA and GLASS cohorts.

Table S4. Significantly enriched Reactome pathways for genes differentially expressed in the TG cohort

Table S5. Significantly enriched Reactome pathways for genes differentially expressed in the GLASS cohort.

Table S6. Significantly enriched Reactome pathways for genes differentially expressed in the TCGA cohort.

Table S7. Significantly enriched Reactome pathways for 217 genes which are differentially expressed in TG, TCGA and GLASS cohorts

Table S8. Significantly enriched Reactome pathways for 72 genes which are differentially expressed in TG and GLASS cohorts.

Table S9. Reactome annotations of TG DE genes included in WGCNA modules according to Sojka et al.

Table S10. Differentially expressed lncRNAs in the TG cohort including their correlating DE targets.

Table S11. Copy Number Alterations of progression-related lncRNAs in the TCGA cohort.

Table S12. Copy Number Alterations of progression-related lncRNAs in the GLASS cohort.

Table S13. Enriched Reactome pathways for the correlating DE lncRNA targets

Table S14. DESeq2 output, comparing miRNA expression profiles between grade 4 vs. grade 2-3 tumors in the TG cohort

Table S15. Differentially expressed miRNAs in the TG cohort including their negatively correlating DE targets.

Table S16. Differentially expressed miRNAs in the TCGA cohort when comparing primary grade 4 vs. primary grade 2-3 tumors.

Table S17. Copy Number Alterations of progression-related miRNAs in the TCGA cohort.

Table S18. Copy Number Alterations of progression-related miRNAs in the GLASS cohort.

Table S19. Enriched Reactome Pathways for the DE miRNA targets

Table S20. Interactions included in the ncRNA-focused Gene Regulatory Network

Table S21. Genes in the ncRNA network that were assigned to a different Reactome pathway category, including references to the supporting literature.

## Statements and Declarations

### Author contributions

A.H.: computational and statistical analysis, result interpretation, manuscript figures, manuscript writing

S.J.: computational and statistical analysis M.A.: computational analysis

J.H., H.H.: sample and clinical data collection, clinical expertise W.Z.: original design of study, supervision

M.N.: data management, supervision

X.L.: design of the gene regulatory network analysis, supervision

K.J.R.: design of study, sample handling, result interpretation, supervision, project coordination

All the authors revised and approved the final manuscript.

### Data availability

Normalized mRNA expression counts are available in GEO with an accession number <u>GSE233034</u>. Normalized miRNA expression counts will be made available under the same GEO series. The raw data from datasets generated during the current study are not publicly available due to lack of patient consent that separately gives permission to this, but the corresponding author can be contacted upon reasonable request. Illumina HiSeq RNA and miRNA gene expression counts for the TCGA-GBM and TCGA-LGG projects used in this study can be retrieved directly from the public GDC data portal (https://portal.gdc.cancer.gov/) or using the R library (https://bioconductor.org/packages/release/bioc/html/TCGAbiolinks.html). Copy number alterations in TCGA-GBM and TCGA-LGG cohorts can be accessed through cBioPortal (https://www.cbioportal.org/study/summary?id=lgggbm_tcga_pub). The gene expression counts and copy number alteration counts from the GLASS cohort can be obtained from the synapse.org portal (https://www.synapse.org/Synapse:syn17038081/wiki/585622). The RNA-data generated by Sojka et al can be obtained from <u>GSE276176</u>.

### Code availability

The code used to generate the figures and results can be found from: https://github.com/CRI-group/ncRNA_IDHmut_astro_progression.

### Ethical approval

The study was performed in line with the principles of the Declaration of Helsinki. Approval for this study was granted by The Regional Ethics Committee of Tampere University Hospital (Tays) (decision R07042, dates 9.2017 and 12.2024) and Valvira (decision V/78697/2017).

### Consent to participate

The requirement for informed consent was waived by The Regional Ethics Committee of Tampere University Hospital (Tays) because of the retrospective nature of the study.

### Consent to publish

Not applicable.

## Acknowledgements

We would like to acknowledge Mrs. Paula Kosonen, Mrs. Päivi Martikainen, Mrs. Marika Vähä-Jaakkola, Mrs. Marja Pirinen, Mrs. Sari Toivola, and Mrs. Hanna Selin for sample handling and logistics. We would like to acknowledge Mr Pauli Helen for his valuable contribution. Personnel at Tampere University Hospital and Fimlab laboratories Ltd. are acknowledged for their contribution to sample collection and M.D. Antti Hyart for frozen sample cohort inventory. We are grateful for them and the patients for permitting the analysis of precious patient material.

The results published here are in part based upon data generated by The Cancer Genome Atlas (TCGA) project (https://www.cancer.gov/tcga) established by the National Cancer Institute (NCI) and the National Human Genome Research Institute (NHGRI). We also acknowledge the valuable contribution of the Glioma Longitudinal AnalySiS (GLASS) data resource to this paper. We acknowledge the CSC - IT Centre for Science, Finland, for providing computational resources.

