## Supplementary figures and images for "Integrative multi-modal analysis reveals the contribution of noncoding RNAs to post-treatment progression of IDH-mutant astrocytomas"

### Supplementary Figure 2

#### Supplementary Figure 2

a)

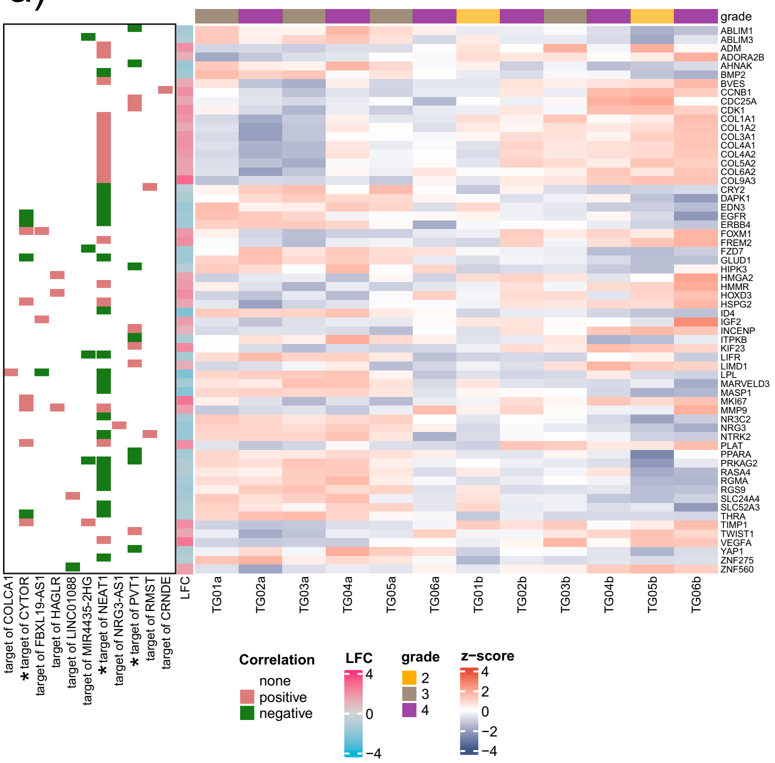


b)

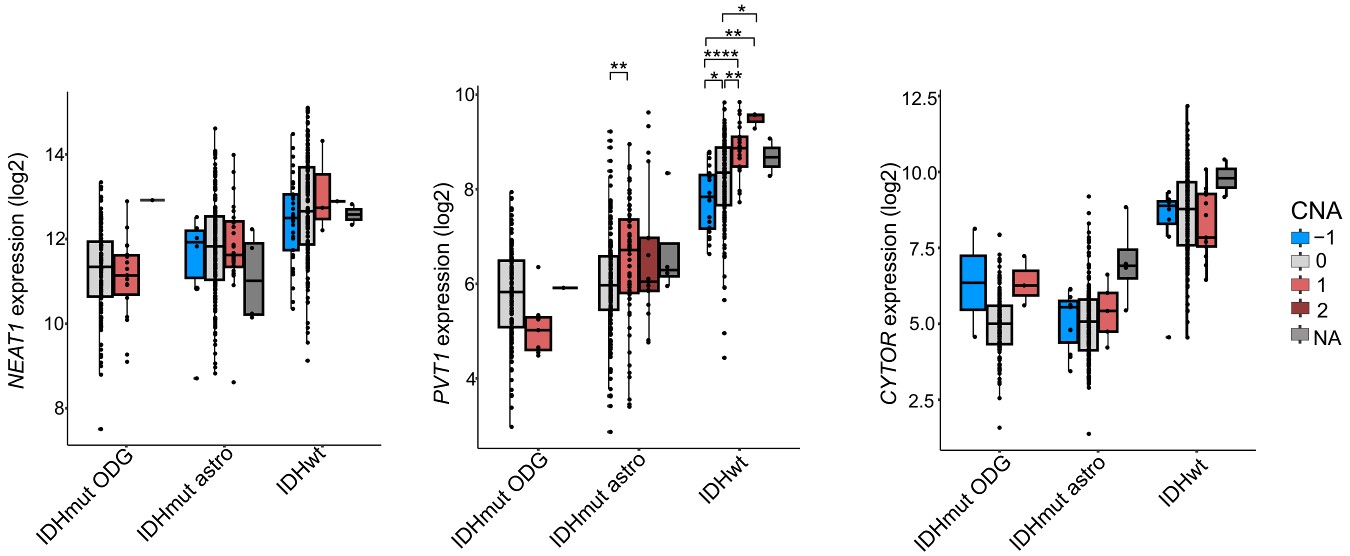

### Supplementary Figure 3

#### Supplementary Figure 3

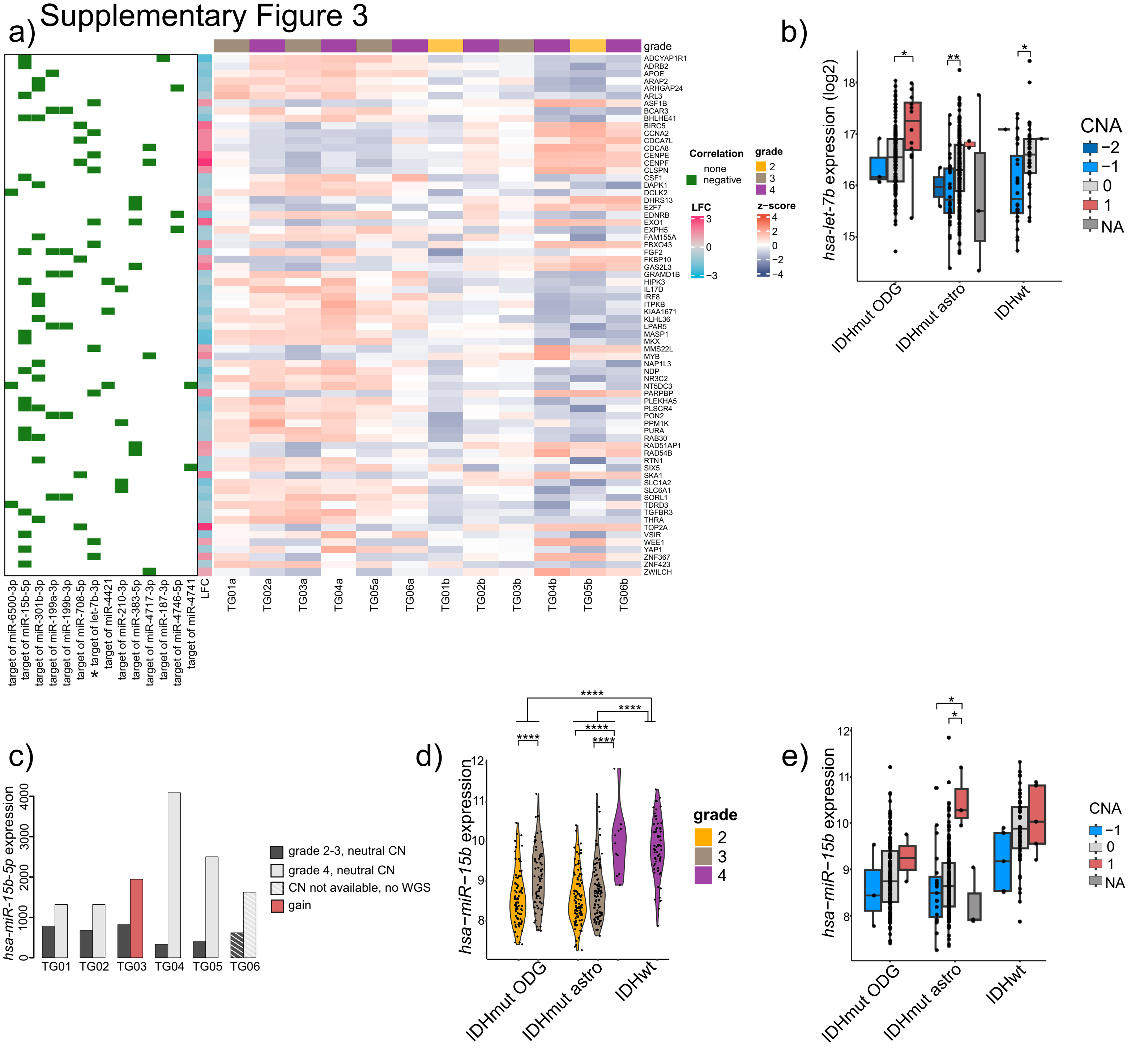

### Supplementary Figure 4

#### Supplementary Figure 4

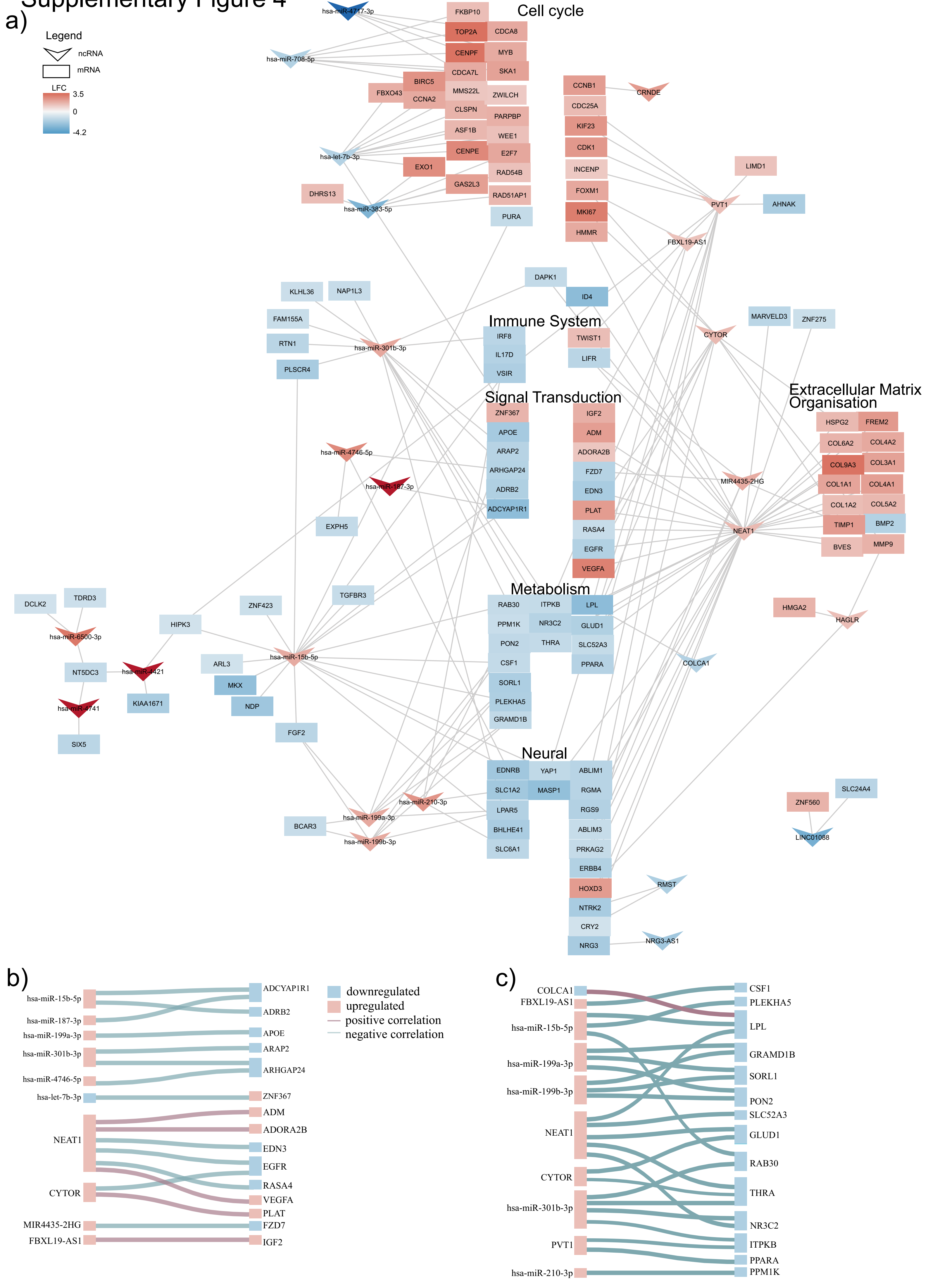
