## Supplementary Figure 1 for "Integrative multi-modal analysis reveals the contribution of noncoding RNAs to post-treatment progression of IDH-mutant astrocytomas"

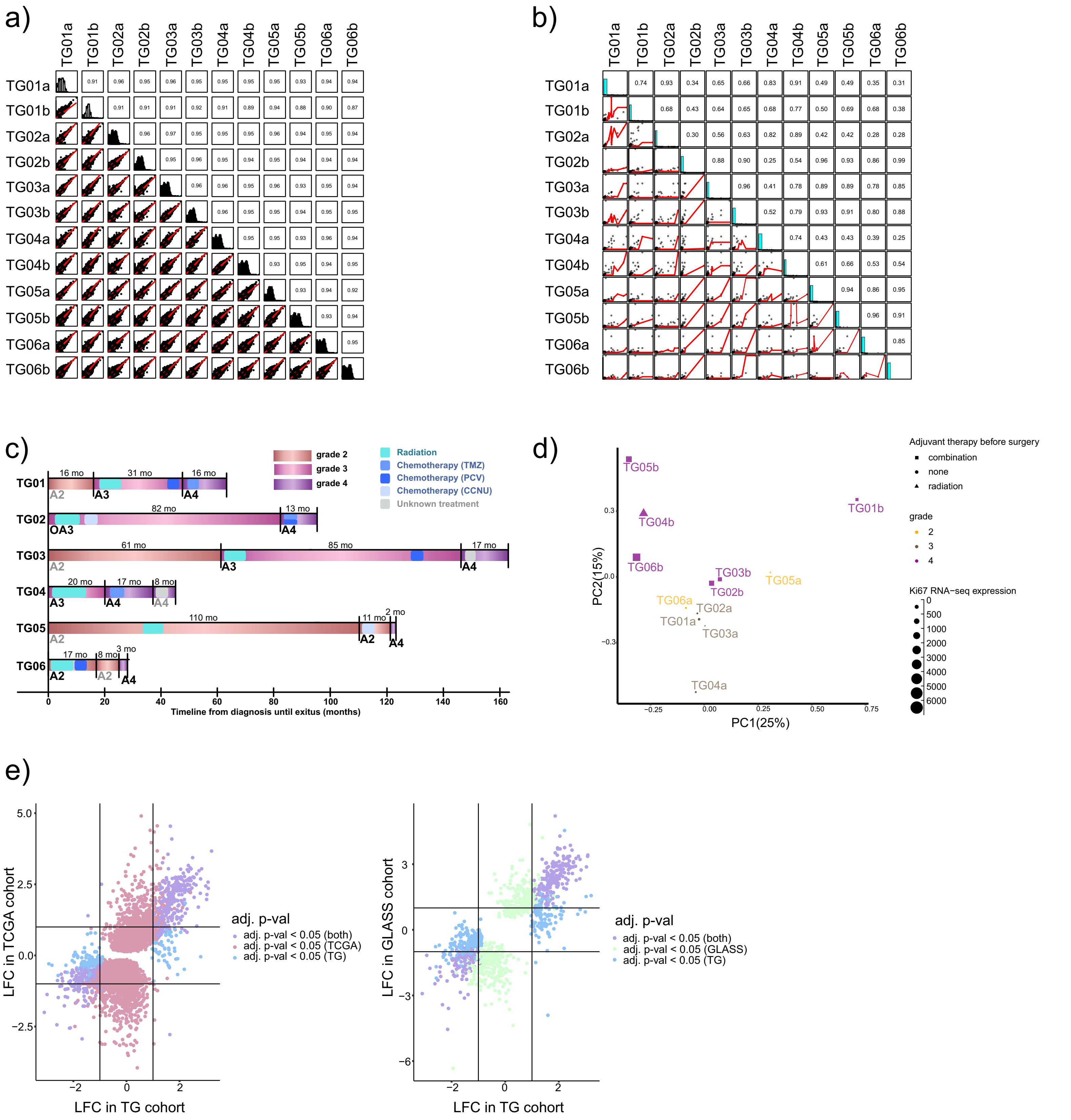

f)

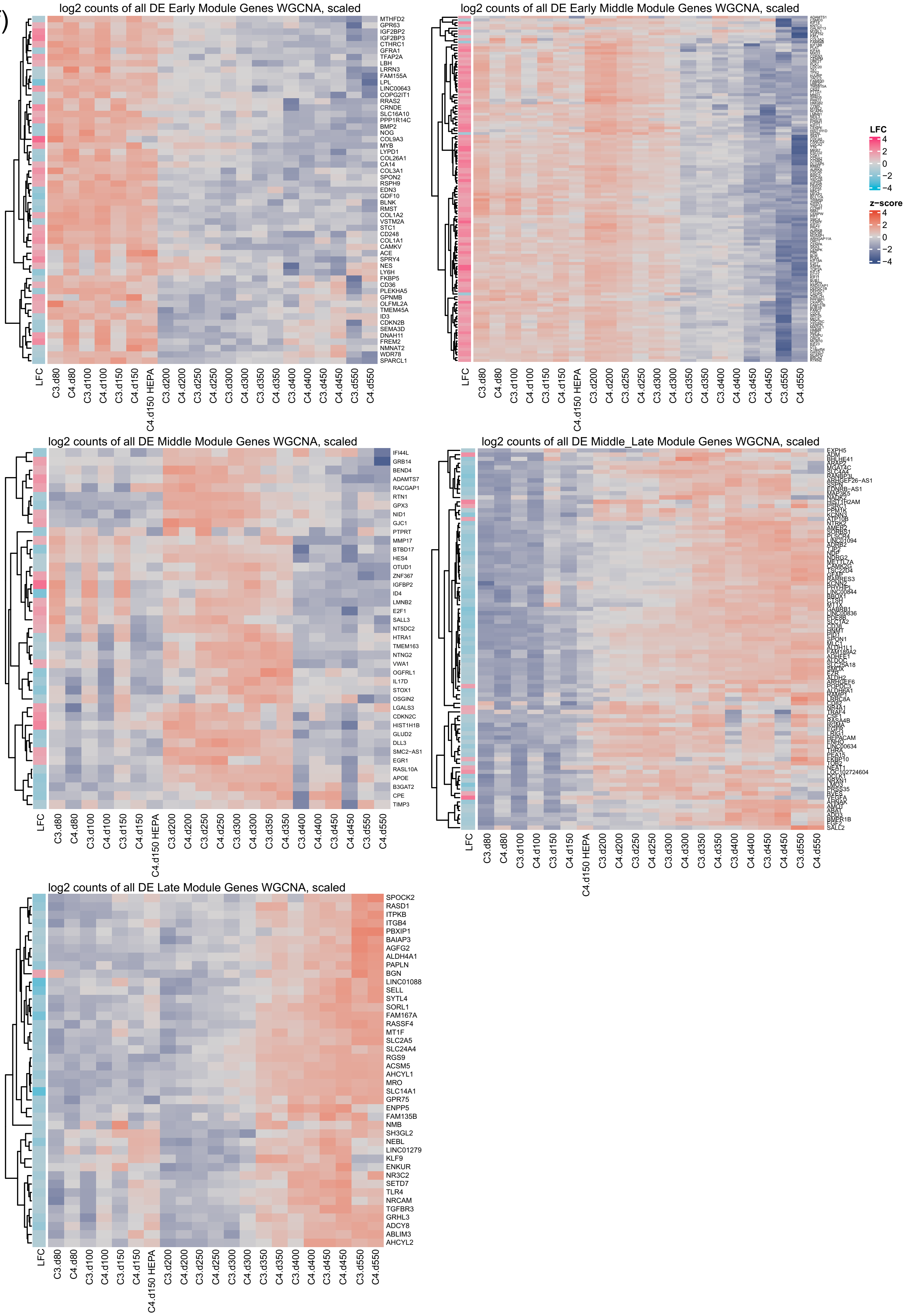
log2 counts of all DE Early Module Genes WGCNA, scaled
MTHFD2
GPR63
IGF2BP2
IGF2BP3
CTHRC1
GFRA1
TFAP2A
LBH
LRRN3
FAM155A
LPL
LINC00643
COPG2IT1
RRAS2
CRNDE
SLC16A10
PPP1R14C
BMP2
NOG
COL9A3
MYB
LYPD1
COL26A1
CA14
COL3A1
SPON2
RSPH9
EDN3
GDF10
BLNK
RMST
COL1A2
VSTM2A
STC1
CD248
COL1A1
CAMKV
ACE
SPRY4
NES
LY6H
FKBP5
CD36
PLEKHA5
GPNMB
OLFML2A
TMEM45A
ID3
CDKN2B
SEMA3D
DNAH11
FREM2
NMNAT2
WDR78
SPARCL1
LFC
C3.d80
C4.d80
C3.d100
C4.d100
C3.d150
C4.d150
C4.d150 HEPA
C3.d200
C4.d200
C3.d250
C4.d250
C3.d300
C4.d300
C3.d350
C4.d350
C3.d400
C4.d400
C3.d450
C4.d450
C3.d550
C4.d550
log2 counts of all DE Early Middle Module Genes WGCNA, scaled
LFC
4
2
0
−2
−4
z−score
4
2
0
−2
−4
log2 counts of all DE Middle Module Genes WGCNA, scaled
IFI44L
GRB14
BEND4
ADAMTS7
RACGAP1
RTN1
GPX3
NID1
GJC1
PTPRT
MMP17
BTBD17
HES4
OTUD1
ZNF367
IGFBP2
ID4
LMNB2
E2F1
SALL3
NT5DC2
HTRA1
TMEM163
NTNG2
VWA1
OGFRL1
IL17D
STOX1
OSGIN2
LGALS3
CDKN2C
HIST1H1B
GLUD2
DLL3
SMC2−AS1
EGR1
RASL10A
APOE
B3GAT2
CPE
TIMP3
log2 counts of all DE Middle_Late Module Genes WGCNA, scaled
log2 counts of all DE Late Module Genes WGCNA, scaled
SPOCK2
RASD1
ITPKB
ITGB4
PBXIP1
BAIAP3
AGFG2
ALDH4A1
PAPLN
BGN
LINC01088
SELL
SYTL4
SORL1
FAM167A
RASSF4
MT1F
SLC2A5
SLC24A4
RGS9
ACSM5
AHCYL1
MRO
SLC14A1
GPR75
ENPP5
FAM135B
NMB
SH3GL2
NEBL
LINC01279
KLF9
ENKUR
NR3C2
SETD7
TLR4
NRCAM
TGFBR3
GRHL3
ADCY8
ABLIM3
AHCYL2
